# Neural response attenuation for shorter inter-onset intervals between sounds in a natural soundscape

**DOI:** 10.1101/2025.05.22.655447

**Authors:** Thorge Haupt, Marc Rosenkranz, Martin G. Bleichner

## Abstract

Sensory attenuation of auditory evoked potentials (AEPs), particularly N1 and P2 components, has been widely demonstrated in response to simple, repetitive stimuli sequences of isolated synthetic sounds. It remains unclear, however, whether these effects generalize to complex soundscapes where temporal and acoustic features vary more broadly and dynamically. In this study, we investigated whether the inter-onset interval (IOI), the time between successive sound events, modulates AEP amplitudes in a complex auditory scene. We derived acoustic onsets from a naturalistic soundscape and applied temporal response function (TRF) analysis to EEG data recorded from normal hearing listeners. Our results showed that shorter IOIs are associated with attenuated N1 and P2 amplitudes, replicating classical adaptation effects in a naturalistic soundscape. These effects remained stable when controlling for other acoustic features such as intensity and envelope sharpness and across different TRF model specifications. Integrating IOI information into predictive modelling revealed that neural dynamics were captured more effectively than simpler onset models when training data were matched. These findings highlight the brain’s sensitivity to temporal structure even in highly variable auditory environments, and show that classical lab findings generalize to naturalistic soundscapes. Our results underscore the need to include temporal features alongside acoustic ones in models of real-world auditory processing.

**Highlights:**

- Neural responses (N1, P2) to sound events are attenuated when inter-onset intervals are short, replicating classic attenuation effects in a naturalistic soundscape.
- Automatic onset detection from complex, ecologically valid soundscapes enables fine-grained analysis of temporal auditory dynamics.
- These findings highlight that temporal sensitivity in auditory processing persists even in highly variable, real-world acoustic environments.

## 1 Introduction

Non-invasive neuroimaging tools, such as electroencephalography (EEG), have been invaluable in unraveling the neural underpinnings of auditory perception (Alain and Winkler, 2012; Gutschalk and Dykstra, 2014; Lee et al., 2014; Rahman et al., 2020). Many of the mechanisms uncovered using EEG have relied on highly controlled, low-complexity stimuli (Crosse et al., 2021; Schutz and Gillard, 2020). Often, unnatural and repetitive, click-like tones have been used, reducing experimental investigation to changes along a single stimulus dimension, such as intensity (Ĺopez-Caballero et al., 2023), frequency (Herrmann et al., 2013, 2014), or inter-stimulus interval (ISI) (Zacharias et al., 2012). An emerging question is whether established neural mechanisms, such as the attenuation of auditory evoked potentials (AEP), can be applied to understand human perception in response to complex and naturalistic soundscapes, where many of the investigated factors occur simultaneously.

A well-documented finding is that the amplitude and latency of an AEP in response to a sound are dependent on the characteristics of the preceding sound, as well as the context. Studies have shown that repeated presentations of tones modulate AEP components associated with acoustic processing, particularly the N1. It has been shown that the N1 amplitude is reduced by a preceding tone for up to 10 seconds (López-Caballero et al., 2023; Wang et al., 2008). Vital to the understanding of the mechanism has been the work by Zacharias et al. (2012). They demonstrated that in random presentation designs, both the interval to the last stimulus and the broader presentation history significantly impact neural response attenuation. In particular, they have shown that the N1 amplitude is modulated in a nonlinear fashion. While the peak modulation has been shown reliably, the exact neural mechanisms driving this attenuation remain debated (May and Tiitinen, 2010; Näätänen and Picton, 1987). Taken together, these findings demonstrate that the N1 component is sensitive to specific stimulus properties, such as temporal spacing. Changes in these acoustic properties lead to predictable patterns of modulation in amplitude and latency of the neural response.

Recent advances in data analysis and experimental designs have made it feasible to study auditory processing in response to more naturalistic sound environments (Brodbeck et al., 2023; Crosse et al., 2021; Holdgraf et al., 2017; Lalor et al., 2009) and situations (Ladouce et al., 2021; Rosenkranz et al., 2023, 2024). The trend towards using more naturalistic stimuli is particularly notable in language research, where studies have gradually moved from presenting isolated words and phonemes (Lutzenberger et al., 1994; Näätänen, 2001) to sentences (Desai et al., 2021), continuous speech (Ding and Simon, 2014; Howard and Poeppel, 2010), and, ultimately, naturally recorded speech (Agmon et al., 2023). This shift towards naturalistic stimuli provides new insights into auditory processing and raises the question of how far results based on experiments using isolated tones generalize to real-world soundscapes (Hamilton et al., 2021; Schutz and Gillard, 2020; Vallet and van Wassenhove, 2023).

Temporal response functions allow the study of neural responses to continuous acoustic stimuli (Crosse et al., 2021; Holdgraf et al., 2017; Kriegeskorte and Douglas, 2019). This approach provides an opportunity to investigate whether our understanding of auditory mechanisms based on isolated tones can be extended to complex, real-life auditory environments. The benefit of these models is that they are straightforward to interpret and allow for the comparison of multiple models (Crosse et al., 2016). However, many standard TRF implementations assume, either explicitly or implicitly, that neural responses to repeated instances of a given feature type, such as peaks in the speech envelope, are uniform. This assumption oversimplifies neural dynamics, as accumulating evidence suggests that brain responses to acoustic features are often nonlinear and context-dependent (Buzsáki and Mizuseki, 2014; Herrmann et al., 2016; Stam, 2005; Wang et al., 2008). This raises a critical question: How can prediction-based models account for the non-linear and temporally dynamic nature of auditory processing?

One such attempt to generalize lab findings to naturalistic soundscapes and account for non-linear dynamics of the brain has been done by Drennan and Lalor (2019). Instead of using a single neural model to describe the speech stimulus, they subdivided their feature, the acoustic envelope, based on soundscape intensity (amplitude value). By accounting for the non-linear neural response to differences in the intensity of the continuous envelope, they developed a more accurate model to explain the neural underpinnings of naturalistic soundscape perception.

Here, we build on the approach of Drennan and Lalor (2019) and extend it to a complex, naturalistic soundscape that is dominated by non-speech sounds. The investigation of the effect of inter-onset interval (IOI) (i.e., the temporal distance between two onsets) on neural response amplitude is non-trivial, since sound events rarely occur at a steady rhythm and differ widely in their acoustic properties. We aim to investigate whether the influence of IOI on neural response amplitude, which was previously observed using simple, isolated stimuli, can be extended to complex, naturalistic soundscapes. Generalizing this relationship to real-world auditory input would provide a framework for understanding brain dynamics in a more ecologically valid setting. Specifically, we test whether the amplitude of the neural response to a sound onset depends on the duration of the interval preceding it, even in the presence of continuous, naturalistic auditory input.

## 2 Method

### 2.1 Data Set

The current study uses an existing data set by Rosenkranz et al. (2023), where they investigated the effect of attentional modulation on auditory perception during a complex audio-visual motor task. Specifically, the soundscape was created to simulate sounds encountered in an operating room to determine the neural response to different types of relevant and irrelevant sounds depending on the attentional instructions.

### 2.2 Task

Participants had to perform a complex visual-motor task that comprised playing 3-dimensional Tetris. In addition to the standard Tetris rules, vocal instructions occasionally told the participants specific locations where to place the blocks. Besides the vocal instructions, participants had to respond to tones. There were two conditions in which the participants had to respond to different tones. They are, however, not relevant for the current analysis.

### 2.3 Soundscape

The soundscape was designed to mimic an operating room, specifically geared towards a surgeon’s perspective. The acoustic environment consisted of speech sounds and environmental sounds (e.g., clattering of tools, footsteps, conversations). The sounds were either vocal instructions on where to place the next block or conversation snippets from a podcast. In total, each participant had to comply with the vocal instructions 12 times, where they were told to “Place the next stone in the [upper\lower left \right] corner”. Importantly, the instructions were played randomly and never consecutively repeated. The conversation snippets were taken from a podcast and also placed randomly, but in semantically coherent order. The content of the conversation snippets was irrelevant to the experiment. In total, 48 snippets were played and lasted roughly 3.5(±1.5)s. The total soundscape was played for roughly 16 minutes per condition, totaling 32 minutes of recorded data on average.

Besides speech segments, the soundscape also contained hospital sounds of people moving around and air conditioning. Furthermore, there were three different tones inserted: alarm, beep, and irrelevant. The alarm and irrelevant sound were 200ms, and the beep tone was 60ms long. Each tone was played 48 times and was also randomly placed into the soundscape. Importantly, the alarm tone was always played from the same direction, whereas the beep was played from multiple directions.

All sounds included in the soundscape were processed in MATLAB, such that the root-mean-squared (RMS) was consistent across them. Accounting for differences in loudness was done by adjusting the loudness through sound-specific gain parameters. Lastly, using the head-related impulse function, tones were spatially separated. The experimental audio stimuli were sampled at 44.1 kHz. All recorded data streams were synchronized via the Lab Recorder software, utilizing the Lab Streaming Layer for integration. Participants provided informed consent after being briefed on the procedure. For more details, see the original paper (Rosenkranz et al., 2023).

### 2.4 EEG Measurement

Participants were fitted with 24 Ag/AgCl passive electrodes positioned according to the 10-20 international system (EasyCap GmbH, Hersching, Germany) for EEG recording. Data collection was performed using a wireless SMARTING system (mBrainTrain, Belgrade, Serbia), with signals referenced to Fz and grounded to AFz. Sampling occurred at a frequency of 500 Hz, and electrode impedance was kept below 20 Ω prior to recording.

### 2.5 Preprocessing of EEG Data

EEG preprocessing was conducted using MATLAB (version 2021a, MathWorks, Natick, MA) with the EEGLab plugin and supplementary custom scripts. Artifact detection was performed using Independent Component Analysis (ICA). To optimize ICA weight estimation, separate preprocessing steps were employed, as recommended by Winkler et al. (2015). This preprocessing pipeline was solely designed for ICA computation and thus did not influence the data ultimately used for analysis. After deriving the ICA weights, they were applied to the unprocessed raw data.

Initially, data from both experimental conditions were combined for each participant. The combined data was resampled to 250 Hz and subjected to a series of filters, starting with a high-pass filter (cutoff: 1 Hz, order: 568) followed by a low-pass filter (cutoff: 42 Hz, order: 128). These cutoff frequencies were chosen to mitigate drifts and line noise, facilitating optimal ICA weight estimation (Winkler et al., 2015). Channels exhibiting poor signal quality were removed using the clean channels function. The data was segmented into 1-second epochs, converted to double-precision format, and artifactual trials were removed using the pop jointprob function with a threshold of three standard deviations.

ICA was executed using the pop runica function with the extended ICA algorithm. The resulting ICA weights were then reapplied to the raw, unfiltered data from each experimental condition. Automatic classification of ICA components as muscle, eye, heart, line noise, or channel noise artifacts was performed using the pop icaflag function, with a predefined probability threshold ([0.7, 1; 0.7, 1; 0.6, 1; 0.7, 1; 0.7, 1]).

Following artifact removal, the raw data underwent a second round of filtering, this time with modified parameters. A low-pass filter was applied first (cutoff: 20 Hz, order: 100), followed by resampling to 100 Hz and high-pass filtering (cutoff: 0.3 Hz, order: 518). The reduced low-pass filter order minimized artifacts associated with steep roll-offs, as recommended by Crosse et al. (2021). The frequency band was restricted to [0.3, 20] Hz, aligning with findings from speech-tracking studies highlighting the dominance of auditory processing in lower frequency ranges (Di Liberto et al., 2015; Crosse et al., 2016). Finally, the data was rereferenced to the mastoids (TP9/TP10).

### 2.6 Temporal Response Function

The neural time series were analyzed using the mTRF toolbox (Crosse et al., 2016) in MATLAB. This toolbox estimates weights that relate neural responses to stimulus features through convolution. The neural response, ***r***(*t, c*), is modeled as the convolution of channel-specific weights (temporal response function), *ω*(*τ, c*), with the stimulus features shifted by a time lag, *τ*, plus a residual term, *ε*(*t, c*):

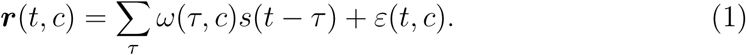

Here, *t* and *c* denote time points and channel indices, respectively. This approach captures the delayed nature of neural responses to stimuli. The resulting weights were analyzed for morphology, topography, model performance, multivariate modeling, and cross-prediction.

The TRF is determined by minimizing the Mean Squared Error (MSE) between observed and predicted neural responses:

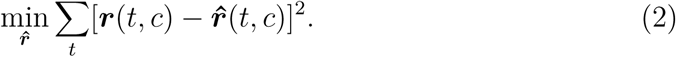

The optimal weights, ***w***, are computed using the formula:

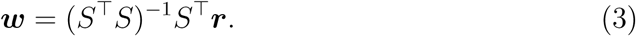

Here, *S* is the design matrix containing the stimulus features across time lags. Its dimensionality is determined by the number of features and lags. Zero-padding was applied at non-zero lags to maintain causality (Mesgarani et al., 2009). The operation *S*^⊤^***r*** represents the inner product between stimulus and neural time series, while (*S*^⊤^*S*)^−1^ accounts for stimulus autocorrelation.

The dimensions of the resulting model weights are determined by the number of features, the time window of integration, and the number of channels. For instance, dividing the soundscape into 8 different bins of IOI intervals yields a model with the dimensionality of 8 × 61 × 22.

### 2.7 Model Training

To train the model, the data was partitioned into 6 segments, where 5 served for training and one segment served as the held-out segment for testing. Within the 5 training segments, cross-validation was applied to derive the optimal lambda for regularization. The resulting model was used to predict the data of the test segment and correlated to the actual neural data. The correlation is the performance marker and is indicative of the prediction accuracy of the model. This approach was consistently applied over all analyses unless stated otherwise. A typical time lag window of [-100, 500] ms was used unless stated otherwise. Cross-validation included a lambda parameter search over values ranging from 10^−4^ to 10^4^ in linear steps of 10.

To investigate the role of training data, we increased the available data and contrasted the bin IOS models against the single onset vector. First, we merged the datasets of the two conditions per participant. We then divided the concatenated data into 12 segments, each serving as a test set once.

### 2.8 Analyses

#### 2.8.1 Features

##### Onsets

We are interested in the effect of the inter-onset interval of two consecutive sounds on the neural response. Unlike previous studies that investigated this effect with pure tones, we are interested here in whether we can replicate the effect in complex soundscapes. For this, we needed to identify sound onsets in the continuous soundscape. To obtain onsets, we used peak detection of acoustic novelty functions (Müller, 2021), which are defined by changes in the energy, spectral flux, and phase changes of the signal, respectively.

##### Energy Novelty

The underlying assumption of the first novelty function is that sound event onset leads to changes in the energy of the signal (*x*). To obtain this representation, the raw signal was squared, and a continuous measure of local energy was obtained by convolving it with a Hann windowing function. Next, the signal was downsampled to the EEG sampling rate at 100 Hz. Given that sound perception of different intensities is logarithmic in humans, we applied a logarithmic compression *log*(1 + *γ* ∗ *x*), where the compression is controlled by *γ*= 10. Finally, the rate of change of the signal was obtained by taking the derivative and half-wave rectifying it.

##### Spectral Novelty

The second novelty function we derived was the spectral flux. Instead of depicting changes in the broadband signal, where overlapping sounds could mask each other, spectral decomposition into different frequency bands can provide a more detailed account of acoustic changes in the signal. First, the signal was decomposed into its frequency components using the short-time Fourier transform (STFT). The magnitude in each frequency band was obtained by taking the absolute value and applying logarithmic compression (*γ*= 10). To determine the rate of change in each frequency band, the first derivative was taken, and the signal was half-wave rectified. At last, the signal was obtained by summing over frequency bands. Postprocessing involved removing small fluctuations by subtracting the local average of the signal. Negative values were set to zero.

##### Complex Novelty

The third novelty function extends the spectral flux function by considering changes in the phase of the signal’s frequency components. To avoid chaotic noise-like phase fluctuations impairing the novelty estimate, the phase is weighted by the magnitude of the Fourier coefficient. That is, phase information becomes only relevant, given the magnitude of the Fourier coefficient. The novelty function was derived by determining the difference between the predicted and actual signal, where larger values refer to greater change. Here, the predicted signal was construed based on the assumption of local stationarity, implying that the phase and magnitude of the Fourier coefficients stay relatively constant over some time.

Similar to the spectral flux, the signal was decomposed into Fourier coefficients using the STFT, and besides the magnitude, phase values were extracted. The angle of the coefficients was derived and normalized by 2*π*. Afterwards, the rate of change was determined by taking the derivative of the phase values. The Fourier coefficient of the next frame was predicted based on the magnitude, current phase, and rate of phase change. The difference between the actual and predicted coefficient was derived, and novelty values smaller than the previous one were set to 0. The novelty function was obtained by summing over frequencies. Local averaging and half-wave rectification were applied to obtain smoother results.

##### Onset Detection

Each novelty function represents a distinct measure of change in the signal, contributing unique information. To leverage the complementary representations of change, we normalized the novelty functions between 0 and 1 and averaged them together. This approach aimed to integrate the advantages of all novelty functions, capturing a more comprehensive depiction of changes in the auditory environment.

To detect sound event onsets, we applied an adaptive thresholding algorithm. Unlike global thresholding, which can overlook smaller, noise-like peaks, adaptive thresholding considers the local temporal structure. Specifically, we smoothed the combined novelty function using a Gaussian window (*σ* = 4) and applied an offset defined as *mean*(*x*) + 0.05. To further refine the signal, we employed a median filter with a window size of 1024 samples. The resulting signal represented the local average, and a peak was only selected if the novelty exceeded the local threshold. The temporal location of each detected peak was recorded as a sound event onset. Finally, the onset information was encoded into a binary feature vector for further analysis (Figure 1).

**Figure 1.**
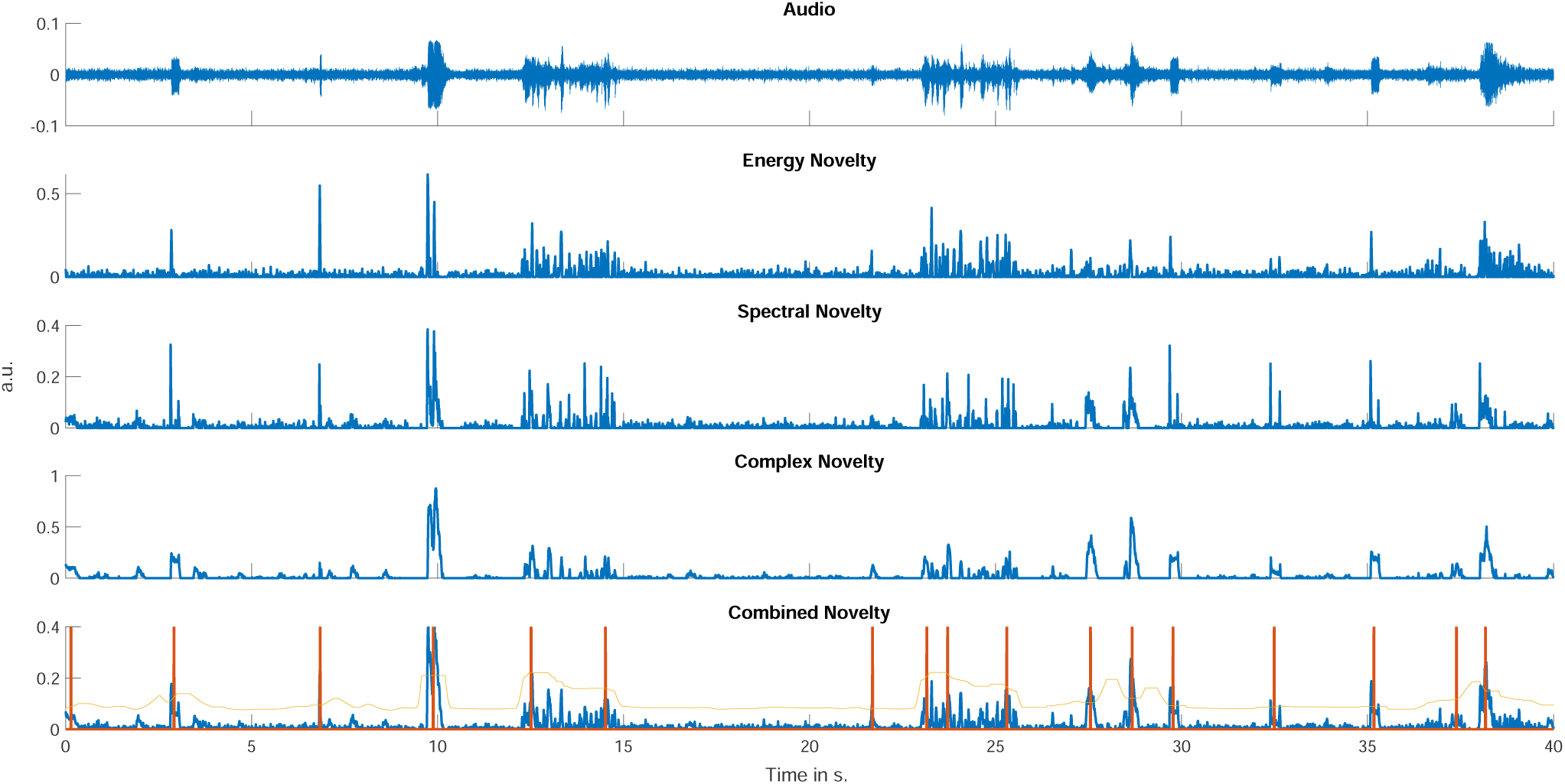
A representation of the novelty functions and the corresponding peak-picking algorithm. The top plot shows the audio signal played to Participant 1 during the narrow condition. Below are the corresponding energy, complex, and spectral novelty functions. The last plot shows the combined novelty functions in blue, the local average in yellow, and detected peaks in orange.

#### 2.8.2 IOI Analysis

For the IOI, we calculated the time interval between successive onsets. Onsets separated by more than 10 seconds were excluded from further analysis, based on previous research findings suggesting that neural attenuation occurs within this time frame (Zacharias et al., 2012).

Inspired by the method of Drennan and Lalor (2019), we applied a similar strategy based on the IOI between sound event onsets. First, we defined ranges for the IOI values and assigned each onset to its corresponding bin. To determine the bin edges, we analyzed the sample distribution of distance values (Figure 2). This distribution of distances of the sound onsets is non-normally distributed and is skewed to the lower distance values. Here, 80% of the sound event onsets follow another sound event within 3.63 seconds. The binning was designed to ensure a uniform distribution of onsets across bins, meaning each bin contained an equal number of onsets. Since no prior studies had applied this approach, we experimented with different bin numbers (ranging from 2 to 8), leading to seven distinct models with varying numbers of bins. These models were then used to predict unseen data, and amplitude values were extracted.

**Figure 2.**
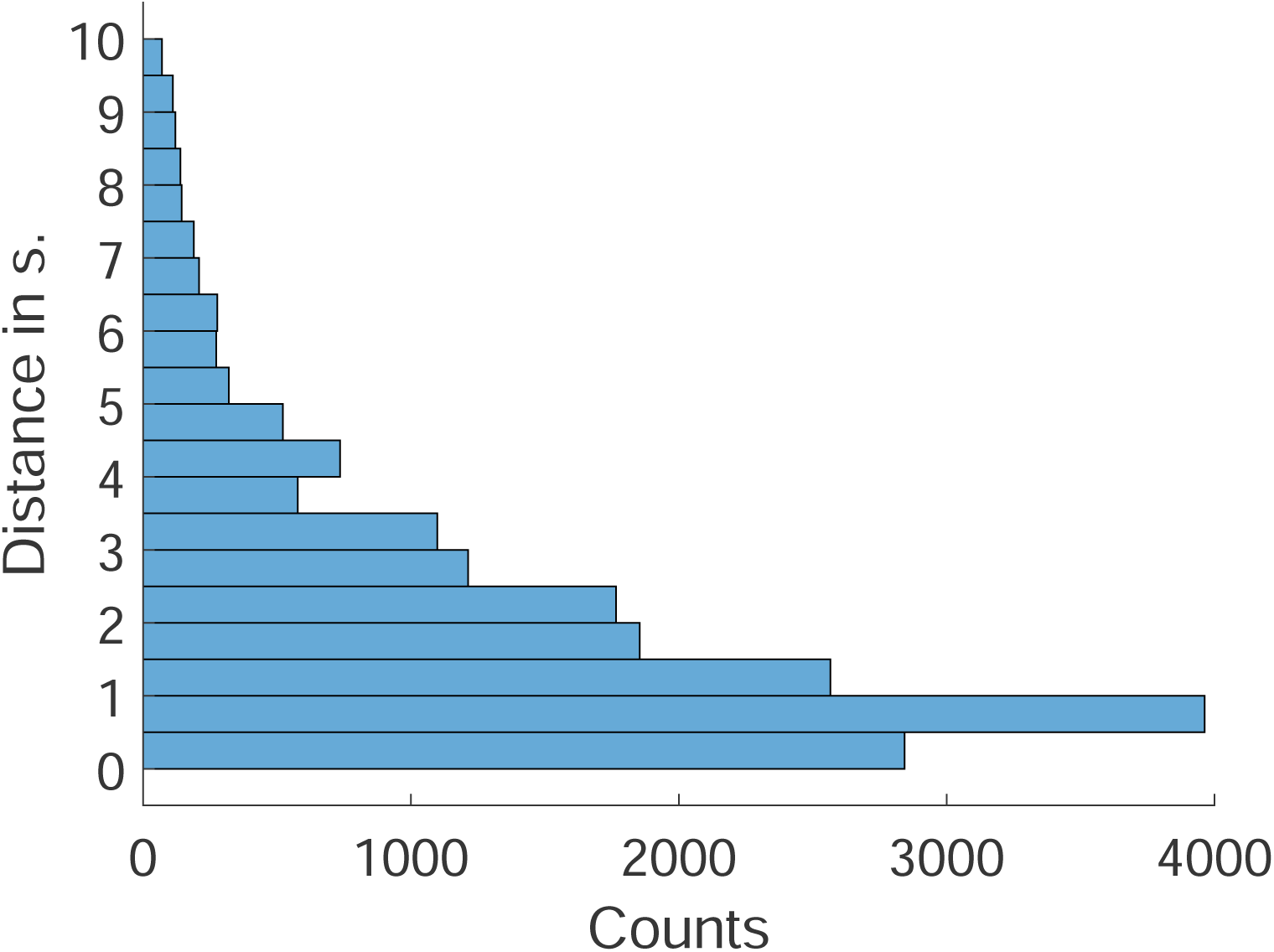
Histogram displaying the distance of onsets to the previous one over all participants. score for comparison.

To determine whether the peak values as a function of IOI were due to chance, we conducted a permutation analysis. Specifically, we randomly shuffled the allocation of sound onsets to their corresponding bins while preserving the overall distribution of onsets. This ensured that changes in the soundscape were still captured, but without a structured relationship to IOI. For each model, we summed the difference between each peak value, representing a rate of change score. The gradient of amplitude values over IOI for each bin model served as the aggregate

We applied the same analysis to 100 permuted model values, generating a distribution of 100 chance gradient values. Statistical significance was asserted at *p <* 0.05. To control for multiple comparisons, we applied a false discovery rate (FDR) correction.

### 2.9 Acoustic Properties Beyond Sound Event Distance

To investigate whether the observed neural amplitude differences between bins could be attributed to systematic acoustic properties of the stimuli, beyond IOI, we extracted two additional acoustic markers known to influence neural response magnitude. The first was the intensity (amplitude) of the sound event. Previous research has demonstrated that sound intensity is positively non-linearly related to neural response amplitude (Adler and Adler, 1989; Drennan and Lalor, 2019; López-Caballero et al., 2023). Interestingly, López-Caballero et al. (2023) examined both intensity and inter-stimulus interval (ISI) and found that both modulated the N1 and P2 components. Moreover, they reported a positive interaction between these factors, specifically, the modulatory effect of intensity on neural responses was more pronounced at longer ISIs, suggesting a dynamic interplay between temporal and intensity cues. The second factor was the sharpness of the envelope onset. This characteristic has posed challenges in auditory research, as sound events with slow-rising envelopes complicate the accurate determination of perceptual onset (Rosenkranz et al., 2024). In such cases, automatic onset detection may not identify the optimal alignment point, potentially resulting in temporal smearing when responses are averaged. To quantify envelope sharpness, we calculated the gradient of the waveform within the first 50 ms following onset.

For each sound event, we derived values for these two markers: amplitude and sharpness, alongside the IOI to the preceding event. To assess their respective contributions, we employed a linear mixed-effects modeling approach. Due to the low signal-to-noise ratio associated with neural responses to rapidly successive events, we did not model single-trial neural response amplitudes directly. Instead, we used IOI as the dependent variable to examine its relationship with the other acoustic predictors.

## 3 Results

Previous research has established that for isolated tones, neural attenuation occurs when tones are played in close succession (Ĺopez-Caballero et al., 2023; Wang et al., 2008; Zacharias et al., 2012). Here, we extend these findings by examining whether a similar modulation occurs in more complex, naturalistic auditory environments. Specifically, we examined whether the neural response to sound events is modulated by the inter-onset interval. To test whether accounting for varying IOI of sound onsets would show modulation of neural response in naturalistic soundscapes, we derived models by grouping sound event onsets in specific IOI ranges. We then tested whether the grouping of sound onsets into varying distance bins would also explain more neural variability.

### 3.1 Modulating Acoustic Properties

To evaluate the potential influence of systematic acoustic differences of the sound events of the different bins, we applied a linear mixed-effects modeling approach, with participants modeled as random intercepts. Before running the model, we assessed collinearity between predictors and found significant correlations between distance and intensity (*r* = −0.15*, p <* 0.001) and between sharpness and intensity (*r* = 0.35*, p <* 0.001), suggesting moderate interdependence among these variables. The correlation could impair the estimated coefficients.

The final model included sharpness, intensity, and their interaction (sharpness ∗ intensity) as fixed effects, with participants as random intercepts. The analysis revealed a significant main effect of intensity (*β* = −195.92*, SE* = 9.75*, t*(18, 327) = −20.09*, p <* 0.001), indicating that higher sound intensity was reliably associated with shorter IOIs. In contrast, the main effect of sharpness was not significant (*β* = 12.64*, SE* = 14.99*, t*(18, 327) = 0.84*, p* = 0.399), nor was the interaction between sharpness and intensity (*β* = 17.41*, SE* = 21.33*, t*(18, 327) = 0.82*, p* = 0.415).

The estimated variance of the random intercept for participants was negligible (4.36∗10^−14^), suggesting minimal inter-individual variability in baseline inter-event distances. These results indicate that intensity is a robust negative predictor of IOI, while sharpness and its interaction with intensity do not contribute significantly to explaining variability in distance.

### 3.2 Peak Modulation

Our results revealed amplitude modulation based on inter-onset interval. The larger the IOI, the larger the amplitude of the AEP. This finding was consistent across all variations of binning parameters. Neither the number of bins nor the specific constraints applied to the binning process significantly altered these findings. Specifically, we observed that the amplitude of the neural response was enhanced for sound events that followed a preceding sound at a greater IOI (Figure 3). To quantify this effect, we extracted peak values of the N1 and P2 components from group-averaged temporal response functions. The results demonstrated a clear trend in which neural response amplitude increased as a function of IOI between tones. Notably, we found that greater temporal spacing elicited a larger N1 peak and a stronger P2 peak across all binning variations (Figure 3).

**Figure 3.**
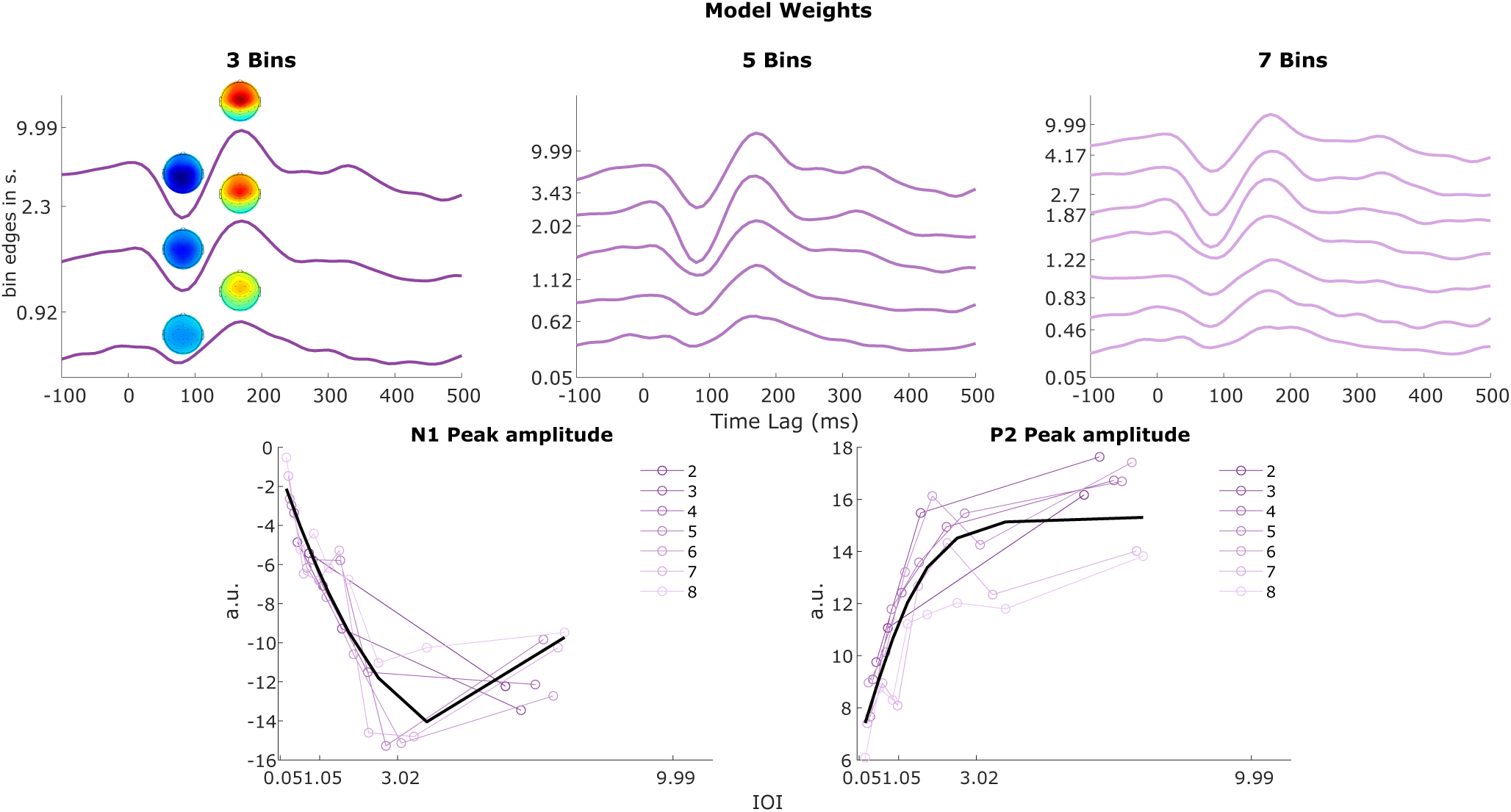
The top row shows model weights of three different bin models, i.e., 3, 5, and 7 bins. For the three-bin model, we also show the topographies at the N1 and P2 latencies for the different model weights. The Y-axis shows the upper edge of each bin. The bottom row shows the minimum and maximum magnitude of the N1 and P2 waves, respectively. The values are extracted for each bin of the seven different models. The models differ in their total number of bins, and bin edges vary to the uniform distribution constraint. On the bottom row, the left plot shows the distribution of N1 peak values as a function of bin edges. On the right, the same is displayed for the P2 values. The black line indicates the optimal model, fitted to all values.

To determine the relationship between the peak amplitudes and IOI, we fit a logistic, exponential, and polyfit model to the data. The first two models were inspired by existing literature (Herrmann et al., 2016; Zacharias et al., 2012). The results showed that the optimal model to describe N1 was the polyfit model of second degree (*R*^2^ = 0.84) and the logistic model for the P2 (*R*^2^ = 0.75).

The results of the permutation testing revealed that the change of amplitude of the N1 and P2 for increasing IOI was significantly above the chance level (*p <* 0.000) (Figure 4). This effect was found for all models.

**Figure 4.**
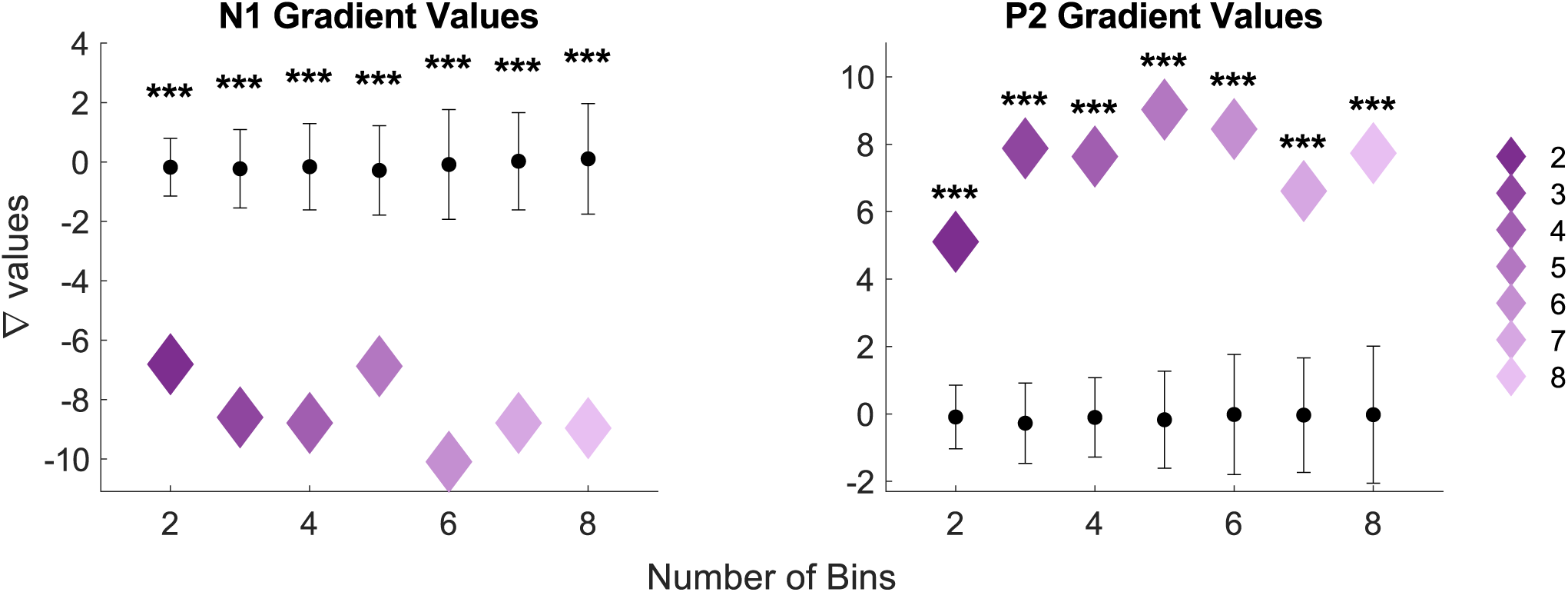
Displayed are the gradient amplitude values of the N1 and P2 for each binned model, respectively. The black bars indicate the standard deviation, and the dots the mean of the chance-level permutation values. The significance level here is indicated at \**p <* 0.05, \*\**p <* 0.01, and \*\*\**p <* 0.000.

### 3.3 Prediction Analysis

Next, we examined whether incorporating IOI information into the neural model estimation improved the explained variability in the recorded signal. To do this, we compared the prediction accuracies of multiple models. First, we tested whether our IOI binned model outperformed chance-level predictions based on the results of the permutation testing (Figure 5). The results showed that all models outperformed the random models significantly: (**2**: *W* = 202, *Z* = −3.62, *p* = 0.002, *ρ* = 0.57; **3**: *W* = 208, *Z* = −3.85, *p* = 0.001, *ρ* = 0.61; **4**: *W* = 204, *Z* = −3.7, *p* = 0.002, *ρ* = 0.58; **5**: *W* = 207, *Z* = −3.81, *p* = 0.001, *ρ* = 0.60; **6**: *W* = 210, *Z* = −3.92, *p* = 0.001, *ρ* = 0.62; **7**: *W* = 202, *Z* = −3.62, *p* = 0.002, *ρ* = 0.57; **8**: *W* = 198, *Z* = −3.47, *p* = 0.003, *ρ* = 0.55). The results indicated a non-linear relationship between model dimensionality and prediction accuracy. Specifically, the difference in prediction accuracy between structured and random models varied depending on the number of bins used. At extreme levels of model dimensionality, either very low or very high, the performance of the binned model only marginally exceeded the chance level. In contrast, intermediate levels of dimensionality produced the most pronounced differences in prediction accuracy. The greatest improvement occurred when using 3-6 bins, suggesting an optimal balance between information preservation and model complexity.

**Figure 5.**
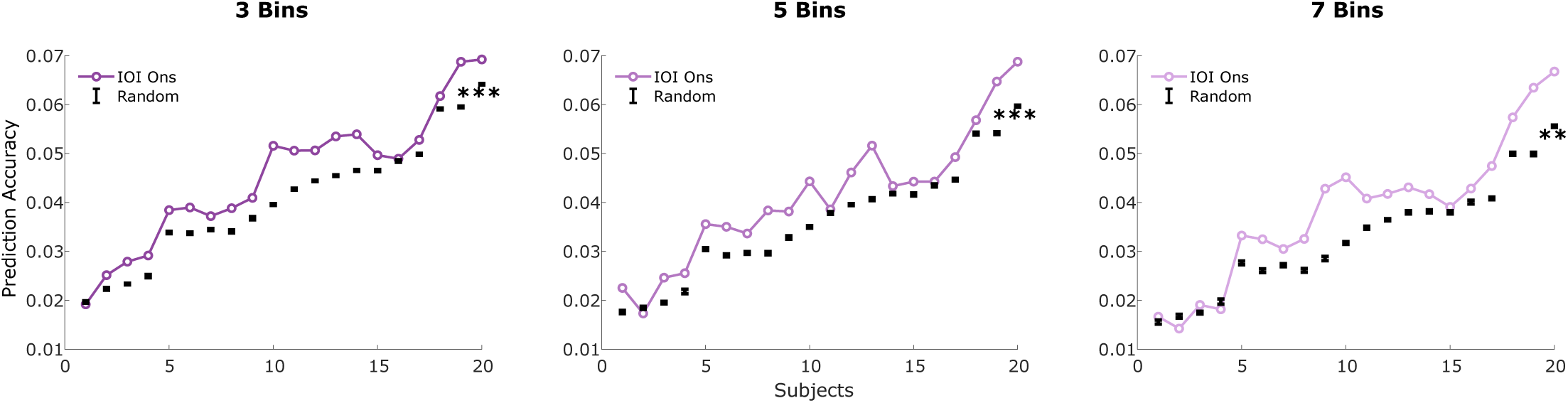
The plot shows the comparison of prediction accuracies between the binned models, i.e., 3, 5, and 7, and the random permutation model over participants. The significance level here is indicated at \**p <* 0.05, \*\**p <* 0.01, and \*\*\**p <* 0.000.

#### 3.3.1 Single Model Comparison

Following our comparison of the binned IOI models to the random models, we aimed to determine whether the inclusion of binned temporal information would outperform a simpler model based on a single binary onset vector. Specifically, we contrasted our more complex model to the single-vector case, which does not incorporate binning information.

Despite observing amplitude modulation as a function of the temporal spacing of sound onsets, incorporating this information into model derivation did not yield higher prediction accuracies compared to using a single onset vector. In fact, the single onset model consistently outperformed the binned models across all cases (**3**: *W* = 176, *Z* = 2.65, *p* = 0.039, *ρ* = 0.42; **4**: *W* = 210, *Z* = 3.92, *p* = 0.001, *ρ* = 0.62; **5**: *W* = 210, *Z* = 3.92, *p* = 0.001, *ρ* = 0.62; **6**: *W* = 208, *Z* = 3.85, *p* = 0.001, *ρ* = 0.61; **7**: *W* = 210, *Z* = 3.92, *p* = 0.001, *ρ* = 0.62; **8**: *W* = 210, *Z* = 3.92, *p* = 0.001, *ρ* = 0.62). The only exception was the simplest binned model, which divided the data into two bins (**2**: *W* = 128, *Z* = 0.86, *p* = 1, *ρ* = 0.14) (Figure 6, middle). These results were stable regardless of whether binning was performed using linear or logarithmic spacing or when sample points per bin were uniformly distributed, as was the case here for the presented results. Additionally, accounting for condition differences did not significantly impact model performance.

**Figure 6.**
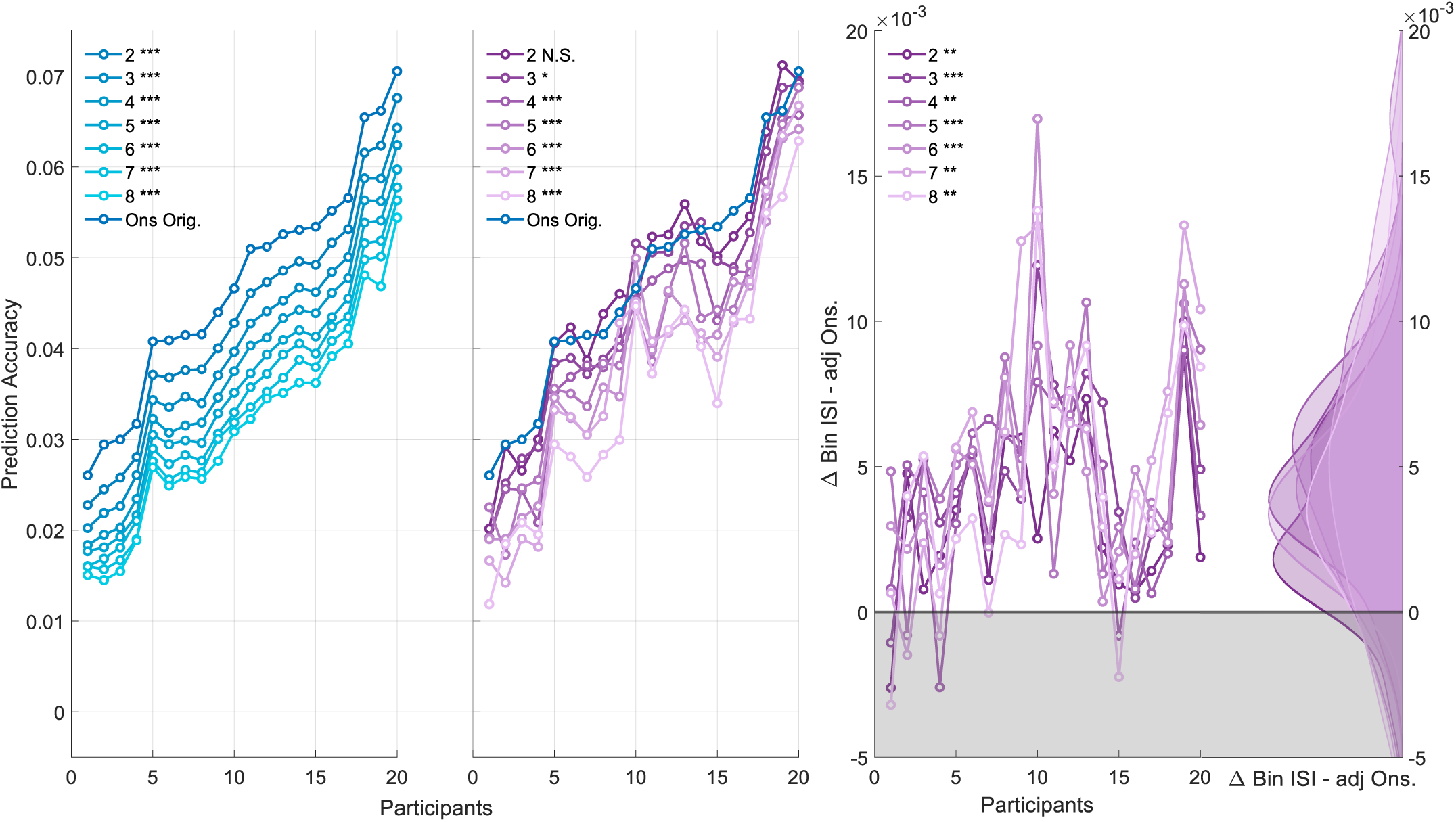
A shows the prediction accuracies over participants for different models. The left plot shows the prediction accuracies for the training adjusted onset model, which is based on the average number of onsets of the corresponding bin IOI model. The plot in the middle contrasts the prediction accuracy of the seven bin IOI models with the onset model. The plot on the right shows the difference between the bin IOI model and adjusted onset.

Furthermore, we observed a deterioration of the prediction accuracy with an increasing number of dimensions. This suggests that data available for training may be a key factor driving the observed difference between the single and binned models. One potential concern is that the onset model and the binned IOI model may not be entirely comparable due to differences in the quantity of training data available for each set of feature weights.

To test whether, at comparable amounts of training data, our bin IOI model would capture neural data more optimally compared to the single onset model, we revisited the previous analysis. Specifically, we adjusted the number of onsets used for training in the single onset vector model to match the average number of onsets per bin for all binned models. For instance, in the case of the 2-bin model, roughly 200 onsets per bin are available. Thus, we randomly selected 200 onsets in the single onset model for training. This process was repeated 100 times for each model, for each participant, and condition.

The results, shown in Figure 6, contrast the performance of the adjusted single onset models with the full single onset vector model. Trivially, every adjusted onset vector was outperformed by the full single onset vector (*W* = 210*, Z* = 3.92*, p* = 0.001*, ρ* = 0.62). Notably, the statistics hold for all comparisons, since parametric tests do not consider the mean difference between distributions. Here, the respective statistics of the effect size, z value, W rank, and p value represent the upper bound.

Given the impact of training data availability, we then revisited the comparison between the binned ISI models and the adjusted single onset vectors. The results indicate that the binned ISI models significantly outperformed their adjusted single onset counterparts across all bin variations (**2**: *W* = 199, *Z* = −3.51, *p* = 0.003, *ρ* = 0.78; **3**: *W* = 208, *Z* = −3.85, *p* = 0.001, *ρ* = 0.86; **4**: *W* = 201, *Z* = −3.58, *p* = 0.002, *ρ* = 0.80; **5**: *W* = 208, *Z* = −3.85, *p* = 0.001, *ρ* = 0.86; **6**: *W* = 210, *Z* = −3.92, *p* = 0.001, *ρ* = 0.88; **7**: *W* = 204, *Z* = −3.7, *p* = 0.002, *ρ* = 0.83; **8**: *W* = 198, *Z* = −3.47, *p* = 0.003, *ρ* = 0.78). These findings highlight the critical role of training data availability in the observed model performances for binary features.

#### 3.3.2 Extended Data Analysis

When contrasting the prediction accuracy of the model containing both conditions with the single onset vector, we found that only the 2-bin and 3-bin models did not differ significantly from the single onset vector (**2**: *W* = 103, *Z* = −0.075, *p* = 1, *ρ* = 0.017; **3**: *W* = 142, *Z* = 1.38, *p* = 0.51, *ρ* = 0.31). For the more complex models (i.e., bin size *>* 3), prediction accuracy was significantly lower compared to the single onset vector (**4**: *W* = 194, *Z* = −3.32, *p* = 0.005, *ρ* = 0.53; **5**: *W* = 190, *Z* = −3.17, *p* = 0.007, *ρ* = 0.50; **6**: *W* = 206, *Z* = −3.77, *p* = 0.001, *ρ* = 0.60; **7**: *W* = 204, *Z* = −3.7, *p* = 0.002, *ρ* = 0.58; **8**: *W* = 209, *Z* = −3.88, *p* = 0.001, *ρ* = 0.61).

We then trained a generic model using all available participant data, leaving one participant out as a held-out test set (Figure 7). This was repeated for every participant once. The results showed that every binned model significantly outperformed the generic single onset model: (**2**: *W* = 3, *Z* = −3.81, *p* = 0.001, *ρ* = 0.60; **3**: *W* = 2, *Z* = −3.85, *p* = 0.001, *ρ* = 0.61; **4**: *W* = 8, *Z* = −3.62, *p* = 0.002, *ρ* = 0.57; **5**: *W* = 1, *Z* = −3.88, *p* = 0.001, *ρ* = 0.61; **6**: *W* = 4, *Z* = −3.77, *p* = 0.001, *ρ* = 0.60; **7**: *W* = 6, *Z* = −3.7, *p* = 0.002, *ρ* = 0.58; **8**: *W* = 7, *Z* = −3.66, *p* = 0.002, *ρ* = 0.58).

**Figure 7.**
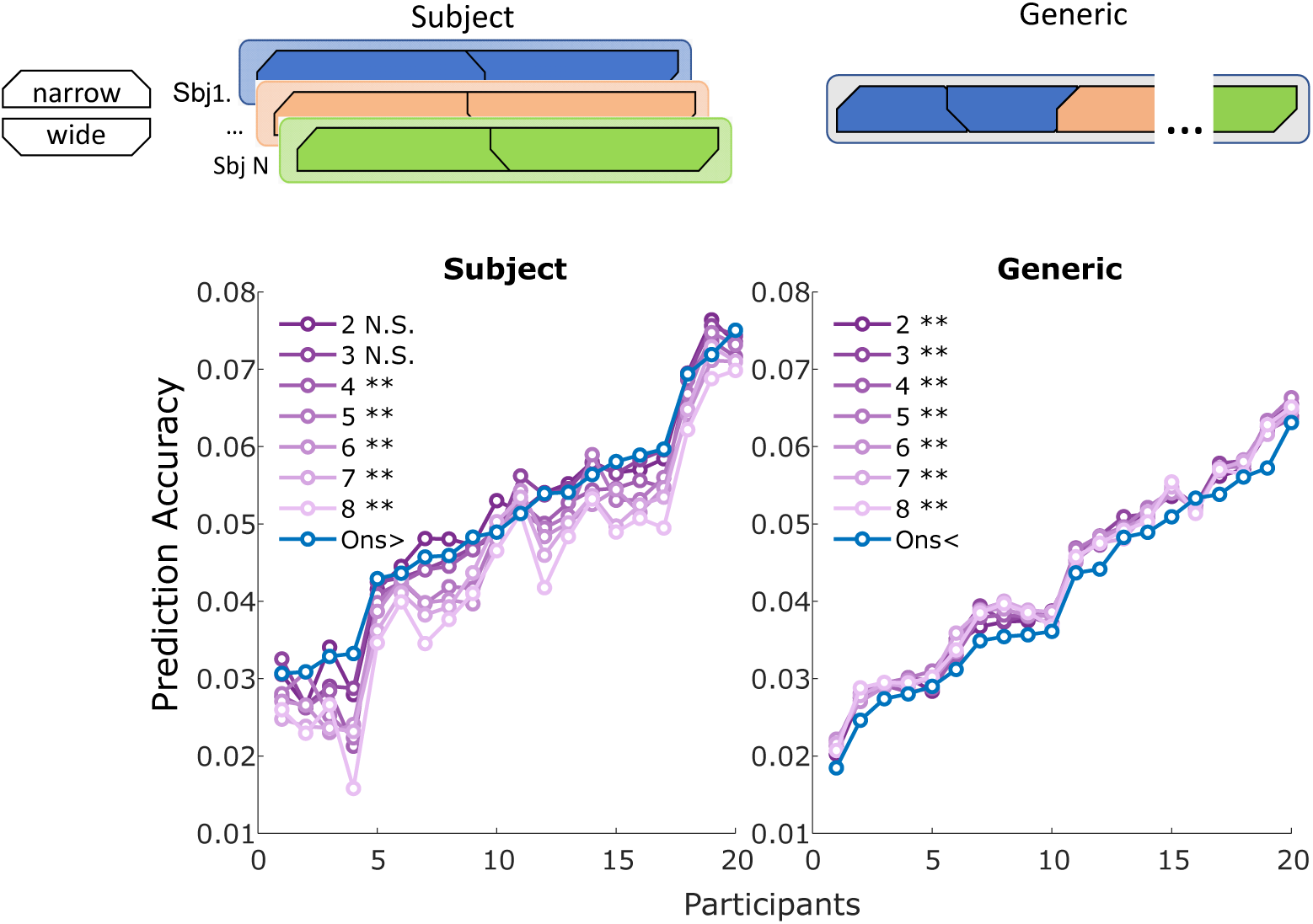
Shows the prediction accuracy of participants for different ways of training the data. The plot on the left shows the prediction accuracy over participants for merged condition data, where a model was trained on this longer dataset. The right side visualizes a generic model being trained. Here, each participant’s merged condition dataset served as a held-out test set once. The *< / >* next to the *Ons* model indicates the direction of significance. The significance level here is indicated at \**p <* 0.05, \*\**p <* 0.01, and \*\*\**p <* 0.000.

#### 3.3.3 Curse of Dimensionality

The relation between training data and training data required is known as the curse of dimensionality, where more complex models require exponentially more training data. One way to mitigate this issue is to parameterize the binary onset vector by the normalized range of distance values. This approach is similar to weighting word onsets in speech processing based on meta-information, such as surprisal. However, adding a parameterized version of the onset vector to the model did not improve prediction accuracy compared to the single onset model (*p* = 1).

## 4 Discussion

Neural response attenuation to acoustic properties has been investigated mostly on isolated pure tones (Beauducel et al., 2000; Herrmann et al., 2016; May and Tiitinen, 2010; Wang et al., 2022, 2008). With this study, we extend those findings to naturalistic soundscapes.

This study provides evidence that neural attenuation is modulated by the IOI of sounds, generalizing validated lab findings to natural soundscapes. Here, the amplitude of the measured neural response was smallest when tones were close together and got larger the further tones were apart. This effect was robust over different numbers of pre-defined bins.

Based on these findings, we implemented the information into neural models to determine whether more neural variability could be explained. In summary, accounting for IOI information improves predictions of neural variability, provided that sufficient training data is available.

### 4.1 Effects of IOI on Neural Data

Our results replicate classic attenuation effects observed for tone sequences. (e.g., click trains or tone bursts) (Costa-Faidella et al., 2011; Herrmann et al., 2016; Lanting et al., 2013; Okamoto et al., 2004; Zacharias et al., 2012; Wang et al., 2008). Crucially, we extend these findings to naturalistic soundscapes, showing that neural responses remain sensitive to inter-event timing despite high variability in acoustic properties. This suggests that neural attenuation with IOI is a fundamental organizing principle in auditory processing.

Interestingly, this amplitude increase appears to plateau for IOIs exceeding three seconds and for more complex models in the case of the N1 peak to decrease again. Beyond these values, additional increases in IOI no longer resulted in further amplitude increase. However, it is important to interpret this plateau cautiously, since specific IOIs were not explicitly manipulated but emerged as a consequence of the binning constraint. Furthermore, due to the naturalistic nature of our stimuli, direct comparisons with studies using strictly controlled IOI intervals are limited. As a result, we cannot draw precise conclusions about the exact time point at which this asymptote occurs. Additionally, separate functions best model the N1 and P2 amplitude values, respectively. This suggests that different mechanisms of attenuation underlie the N1 and P2 and thus possibly reflect separate neural generators (Altmann et al., 2008; Herrmann et al., 2016; López-Caballero et al., 2023).

### 4.2 Neural Mechanisms

Neural attenuation is a fundamental principle of sensory processing, observed across all sensory modalities and at multiple levels of the neural hierarchy, from peripheral receptors to cortical areas. This general phenomenon allows organisms to remain sensitive to new and changing stimuli by dynamically adjusting neural responsiveness in the face of repeated or sustained input (Dean et al., 2005; Hicks and McDermott, 2024; Ulanovsky et al., 2003).

Previous research has proposed two primary frameworks to explain auditory neural attenuation: habituation (often framed within a predictive coding framework) and adaptation. Habituation accounts argue that repeated or predictable stimuli generate sensory expectations, leading to reduced neural responses when those predictions are confirmed and increased responses when they are violated (Costa-Faidella et al., 2011; Nääatänen and Picton, 1987; Ruusuvirta, 2021; Silva et al., 2017; Wang et al., 2008). In contrast, adaptation accounts propose a physiological explanation, suggesting that repeated stimulation leads to reduced neuronal responsiveness due to mechanisms such as synaptic fatigue or depletion, independent of stimulus predictability (Budd et al., 1998; López-Caballero et al., 2023; López-Caballero, 2025; May and Tiitinen, 2010; Rosburg et al., 2022; Rosburg and Mager, 2021).

Our study provides a unique opportunity to discuss these accounts because the naturalistic soundscape we used was highly variable spectrally and temporally. This random nature rules out the formation of stable sensory predictions, making a habituation or predictive coding explanation less likely. Additionally, the soundscape’s broad spectral variability suggests that the attenuation effects we observe are not simply the result of adaptation confined to narrowly tuned, organized auditory neurons. Interestingly, research suggests that temporal and spectral characteristics are adapted independently (Briley and Krumbholz, 2013). Thus, there may be separate neural processes underlying spectral and temporal adaptation.

In line, our findings point toward a general temporal sensitivity in auditory processing. We propose that the observed attenuation reflects broader adaptation mechanisms, such as synaptic depression or slow hyperpolarization, that operate independently of spectral content and are consistent with temporal recovery models described in previous work. The optimal parameters to describe the decay/ recovery and exact models to depict the attenuation are still under debate (Herrmann et al., 2016; Lanting et al., 2013; Regev et al., 2021; Wang et al., 2008; Zacharias et al., 2012).

A promising neural substrate for these effects is the extralemniscal auditory pathway, which has been linked to stimulus-specific adaptation, detection of sudden environmental changes, and supramodal modulation of global brain states (Carbajal and Malmierca, 2018; Shine et al., 2023; Somervail et al., 2021; Willmore and King, 2023; Somervail et al., 2025). Importantly, it lacks the strict tonotopic organization of the lemniscal pathway and has been proposed as the neural substrate of the mismatch negativity (MMN), reflecting an error or novelty signal. Given its broader tuning and role in orienting responses, the extralemniscal system may underlie the non-spectral, time-sensitive attenuation we observed when sound events occurred in close temporal succession.

Despite our promising results, it remains an ongoing challenge to determine the underlying neural mechanisms that are responsible for neural attenuation. Although some theories have been proposed, it remains to be shown whether their integration explains attenuation in response to real-world soundscapes. Future studies should continue to systematically vary both spectral and temporal aspects jointly to determine whether the same or different mechanisms underlie the neural attenuation.

Although unclear where the attenuation occurs, our findings suggest that the temporal sensitivity of the auditory system persists even in complex, unpredictable environments. Importantly, they highlight that attenuation processes extend beyond stimulus-specific mechanisms in the face of dynamic sensory input.

### 4.3 Potential Confounding Factors

Given the complexity of the soundscape under investigation, we examined whether systematic acoustic differences existed between sound events as a function of their IOI. Specifically, we focused on two features previously implicated in modulating neural response amplitudes (Drennan and Lalor, 2019; Ĺopez-Caballero et al., 2023) - sound intensity and envelope sharpness.

Correlational analyses and linear mixed-effects modelling revealed a systematic difference in intensity with IOI, where sound events occurring in close succession tended to have a higher intensity than those spaced further apart. This relationship might be partially biased by the adaptive threshold to select onsets. Here, novelty peaks need to surpass a threshold that is based on the context of the soundscape. Thus, successive onsets may need a larger amplitude to exceed the context-driven threshold, driving the negative relationship between IOI and intensity.

Given the well-established association between increased stimulus intensity and stronger neural responses (Adler and Adler, 1989; Drennan and Lalor, 2019; Beauducel et al., 2000; Ĺopez-Caballero et al., 2023), the fact that closely spaced events were more intense should have amplified rather than diminished their evoked responses. However, our results show the opposite effect: sounds occurring with shorter IOIs elicited attenuated neural responses. This dissociation suggests that intensity differences did not drive the observed IOI-related amplitude modulation, which is in line with findings of López-Caballero et al. (2023), who also found a dissociation between these two factors. On the contrary, intensity-related enhancement may have masked part of the IOI effect, making our findings a conservative estimate of the actual modulation associated with IOI.

### 4.4 Unconsidered Factors Influencing Peak Amplitude Modulation

Our analysis, focused primarily on the IOI as a key modulator of neural responses while accounting for sound intensity and sharpness. However, given the complexity of naturalistic soundscapes, other acoustic and contextual factors may have played a role, which were not systematically investigated in the present study.

One such factor is the duration of the preceding sound. Lanting et al. (2013) reported that longer adapter durations led to greater N1 suppression in a paired-click paradigm, highlighting duration as a potential modulator of adaptation. Although this effect was shown using simple tones, its role in complex soundscapes remains uncertain and warrants further investigation.

Contextual predictability is another critical factor. Previous work has demonstrated that neural attenuation depends on the stimulation history (Zacharias et al., 2012; Herrmann et al., 2016). Notably, reduced attenuation effects are found under random IOI conditions compared to highly predictive sequences. How the general context impacts neural attenuation in autocorrelated soundscapes needs to be investigated by future studies. Evidence from a recent behavioural study suggests that response adaptation to stationary soundscapes occurs faster compared to those with increased spectral variability (Hicks and McDermott, 2024).

Finally, spectral similarity of successive sounds and global soundscape statistics has also been implicated in response attenuation. Herrmann et al. (2013) found stronger adaptation for spectrally similar tones. However, studies using more complex stimuli (e.g., vowels, animal vocalizations) have not observed such effects (Altmann et al., 2008; Silva et al., 2017). Given the broadband nature of our stimuli, the influence of spectral similarity remains ambiguous and was not directly tested here.

Taken together, these findings highlight that while IOI is a critical factor in auditory attenuation, other acoustic dimensions such as duration, spectral content, and stimulus context can also influence peak amplitude modulation. Future work should aim to incorporate these variables into more comprehensive models to better disentangle their individual and interactive contributions to auditory processing in naturalistic environments. As such, it would provide insights into how the brain processes complex soundscapes with greater detail.

### 4.5 Prediction Accuracy

#### 4.5.1 Model Performance

We have shown that integrating IOI provides meaningful information, as shown by the comparison between those models with random onset-bin allocation. Since both models contain onsets at identical time points, but only one assigns onsets based on the IOI between successive sound events, while the other does so randomly, the increased accuracy in the informed model indicates that IOI serves as a meaningful feature for neural prediction.

#### 4.5.2 Trainings Data Availability

When we compared the IOI model against a single onset predictor (i.e., the simple onset model), performance was worse. This result was unexpected, given that the previous analysis showed the model to contain meaningful information. These findings also contrast with the study by Drennan and Lalor (2019), who showed that deriving features that account for the non-linear response of the brain improves model estimation and consequently the amount of neural variability explained.

This discrepancy can be explained by reduced training data per predictor: dividing the onset vector into multiple IOI-based bins leads to fewer events per bin. This impairs the derivation of reliable weights. This observation is crucial in explaining the inferior performance of the bin IOI model to the simple onset model. Given that prediction accuracy is strongly influenced by the availability of training data (Desai et al., 2023; Mesik and Wojtczak, 2023). To verify this interpretation, we controlled for data availability by reducing the simple onset model to match the number of events per bin in the IOI model. Under these conditions, the IOI model outperformed the reduced simple model, confirming that IOI carries predictive value.

We further tested this by increasing the available training data, either by pooling conditions within subjects or by training generic models across participants. In both cases, the IOI model benefited from increased data, often surpassing the simple model. Interestingly, the overall performance of generic models was lower compared to the individual models, and the difference between models was no longer visible. The reduced performance in the generic model compared to subject-specific models is likely due to the latter model better capturing individual nuances. This is in line with previous research showing that when sufficient training data is present, generic models underperform compared to subject-specific models (Mirkovic et al., 2015). Beyond this point, the subject-specific model is superior. The lack of difference between the bin IOI models suggests a ceiling effect of training data, indicating that further data would not yield additional performance gains. This underscores the need to balance feature complexity with the amount of available training data to avoid compromising model performance.

While increasing data availability helped address model complexity, we also examined whether simplifying the model could yield similar results. This approach was inspired by speech processing research, where word onset models are supplemented with parametric word surprisal scores to incorporate additional meaningful information into neural response estimation (Brodbeck et al., 2018). Analogously, we weighted binary onsets by their respective distance to the previous onset. How-ever, this approach did not yield significant improvements in prediction accuracy. Highlighting that the binned approach is capturing non-linear neural dynamics.

#### 4.5.3 Practical implications

These findings highlight a key trade-off: while incorporating temporal context (e.g., inter-onset interval, IOI) improves neural response prediction, increased model complexity requires sufficient training data to avoid performance loss. While shorter lab-based recordings may not provide enough data for models to benefit meaningfully from IOI-based features, longer recordings, particularly those collected in real-world, non-laboratory settings, offer an opportunity to leverage temporal information effectively. As longitudinal, everyday-life recordings (Korte et al., 2025b,a; Rosenkranz et al., 2024; Hölle and Bleichner, 2023; Hölle et al., 2021) become increasingly available, incorporating temporal structure such as IOI may significantly enhance model performance and our ability to predict neural responses in complex, naturalistic contexts.

### 4.6 Conclusion

Our results provide important insights into how the brain processes complex soundscapes in everyday life. We showed that temporal structure, specifically, the timing between sound events, is a critical dimension that modulates neural responses, even in highly variable, naturalistic settings. By demonstrating that shorter IOIs attenuate auditory neural responses and that IOI-based models can outperform simpler onset models (when data availability allows), we highlight the importance of integrating temporal features into the study of auditory scene analysis. These findings lay the groundwork for future research linking neural processing of soundscapes to perceptual, cognitive, and behavioural outcomes, advancing our understanding of how the brain interprets and adapts to the acoustic complexity of the real world.

## Author Contributions

- Thorge Haupt: Conceptualization; Methodology; Writing - original draft; Writing - review & editing
- Marc Rosenkranz: Conceptualization; Writing - review & editing
- Martin G. Bleichner: Conceptualization; Funding acquisition; Writing - review & editing

## Acknowledgements

We would like to thank Manuela Jäger and Silvia Korte for the fruitful discussions throughout the development of the study. This work was funded by the Deutsche Forschungsgemeinschaft (DFG, German Research Foundation) under the Emmy-Noether program - BL 1591/1-1 - Project ID 411333557.

## Potential Declarations

During the preparation of this work, the author(s) used ChatGPT 4o and the free version of ChatGPT (mid 2024) in order to improve language and readability of selected sentences. After using this tool/service, the author(s) reviewed and edited the content as needed and take(s) full responsibility for the content of the publication.

## References

Adler, G. and Adler, J. (1989). Influence of Stimulus Intensity on AEP Components in the 80- to 200-Millisecond Latency Range. Audiology, 28(6):316–324. Publisher: Taylor & Francis eprint: https://www.tandfonline.com/doi/pdf/10.3109/00206098909081638.

Agmon, G., Jaeger, M., Tsarfaty, R., Bleichner, M. G., and Zion Golumbic, E. (2023). “Um…, It’s Really Difficult to… Um… Speak Fluently”: Neural Tracking of Spontaneous Speech. Neurobiology of Language, 4(3):435–454.

Alain, C. and Winkler, I. (2012). Recording Event-Related Brain Potentials: Application to Study Auditory Perception. In Poeppel, D., Overath, T., Popper, A. N., and Fay, R. R., editors, The Human Auditory Cortex, pages 69–96. Springer, New York, NY.

Altmann, C. F., Nakata, H., Noguchi, Y., Inui, K., Hoshiyama, M., Kaneoke, Y., and Kakigi, R. (2008). Temporal Dynamics of Adaptation to Natural Sounds in the Human Auditory Cortex. Cerebral Cortex, 18(6):1350–1360.

Beauducel, A., Debener, S., Brocke, B., and Kayser, J. (2000). On the reliability of augmenting/reducing: Peak amplitudes and principal component analysis of auditory evoked potentials. Journal of Psychophysiology, 14(4):226–240. Place: Germany Publisher: Hogrefe & Huber Publishers.

Briley, P. M. and Krumbholz, K. (2013). The specificity of stimulus-specific adaptation in human auditory cortex increases with repeated exposure to the adapting stimulus. Journal of Neurophysiology, 110(12):2679–2688. Publisher: American Physiological Society.

Brodbeck, C., Das, P., Gillis, M., Kulasingham, J. P., Bhattasali, S., Gaston, P., Resnik, P., and Simon, J. Z. (2023). Eelbrain, a Python toolkit for time-continuous analysis with temporal response functions. eLife, 12:e85012. Publisher: eLife Sciences Publications, Ltd.

Brodbeck, C., Hong, L. E., and Simon, J. Z. (2018). Rapid Transformation from Auditory to Linguistic Representations of Continuous Speech. Current biology: CB, 28(24):3976–3983.e5.

Budd, T., Barry, R. J., Gordon, E., Rennie, C., and Michie, P. (1998). Decrement of the N1 auditory event-related potential with stimulus repetition: habituation vs. refractoriness. International Journal of Psychophysiology, 31(1):51–68.

Buzsáki, G. and Mizuseki, K. (2014). The log-dynamic brain: how skewed distributions affect network operations. Nature Reviews Neuroscience, 15(4):264–278. Number: 4 Publisher: Nature Publishing Group.

Carbajal, G. V. and Malmierca, M. S. (2018). The Neuronal Basis of Predictive Coding Along the Auditory Pathway: From the Subcortical Roots to Cortical Deviance Detection. Trends in Hearing, 22:2331216518784822. Publisher: SAGE Publications Inc.

Costa-Faidella, J., Baldeweg, T., Grimm, S., and Escera, C. (2011). Interactions between “What” and “When” in the Auditory System: Temporal Predictability Enhances Repetition Suppression. Journal of Neuroscience, 31(50):18590–18597. Publisher: Society for Neuroscience Section: Articles.

Crosse, M. J., Di Liberto, G. M., Bednar, A., and Lalor, E. C. (2016). The Multivariate Temporal Response Function (mTRF) Toolbox: A MATLAB Toolbox for Relating Neural Signals to Continuous Stimuli. Frontiers in Human Neuroscience, 10.

Crosse, M. J., Zuk, N. J., Di Liberto, G. M., Nidiffer, A. R., Molholm, S., and Lalor, E. C. (2021). Linear Modeling of Neurophysiological Responses to Speech and Other Continuous Stimuli: Methodological Considerations for Applied Research. Frontiers in Neuroscience, 15.

Dean, I., Harper, N. S., and McAlpine, D. (2005). Neural population coding of sound level adapts to stimulus statistics. Nature Neuroscience, 8(12):1684–1689. Number: 12 Publisher: Nature Publishing Group.

Desai, M., Field, A. M., and Hamilton, L. S. (2023). Dataset size considerations for robust acoustic and phonetic speech encoding models in EEG. Frontiers in Human Neuroscience, 16.

Desai, M., Holder, J., Villarreal, C., Clark, N., Hoang, B., and Hamilton, L. S. (2021). Generalizable EEG Encoding Models with Naturalistic Audiovisual Stimuli. The Journal of Neuroscience, 41(43):8946–8962.

Di Liberto, G. M., O’Sullivan, J. A., and Lalor, E. C. (2015). Low-Frequency Cortical Entrainment to Speech Reflects Phoneme-Level Processing. Current Biology, 25(19):2457–2465.

Ding, N. and Simon, J. Z. (2014). Cortical entrainment to continuous speech: functional roles and interpretations. Frontiers in Human Neuroscience, 8.

Drennan, D. P. and Lalor, E. C. (2019). Cortical Tracking of Complex Sound Envelopes: Modeling the Changes in Response with Intensity. eneuro, 6(3):ENEURO.0082–19.2019.

Gutschalk, A. and Dykstra, A. R. (2014). Functional imaging of auditory scene analysis. Hearing Research, 307:98–110.

Hamilton, L. S., Oganian, Y., Hall, J., and Chang, E. F. (2021). Parallel and distributed encoding of speech across human auditory cortex. Cell, 184(18):4626– 4639.e13.

Herrmann, B., Henry, M. J., Johnsrude, I. S., and Obleser, J. (2016). Altered temporal dynamics of neural adaptation in the aging human auditory cortex. Neurobiology of Aging, 45:10–22.

Herrmann, B., Henry, M. J., Scharinger, M., and Obleser, J. (2013). Auditory filter width affects response magnitude but not frequency specificity in auditory cortex. Hearing Research, 304:128–136.

Herrmann, B., Schlichting, N., and Obleser, J. (2014). Dynamic Range Adaptation to Spectral Stimulus Statistics in Human Auditory Cortex. Journal of Neuroscience, 34(1):327–331. Publisher: Society for Neuroscience Section: Brief Communications.

Hicks, J. M. and McDermott, J. H. (2024). Noise schemas aid hearing in noise. Proceedings of the National Academy of Sciences, 121(47):e2408995121. Publisher: Proceedings of the National Academy of Sciences.

Holdgraf, C. R., Rieger, J. W., Micheli, C., Martin, S., Knight, R. T., and Theunissen, F. E. (2017). Encoding and Decoding Models in Cognitive Electrophysiology. Frontiers in Systems Neuroscience, 11:61.

Howard, M. F. and Poeppel, D. (2010). Discrimination of Speech Stimuli Based on Neuronal Response Phase Patterns Depends on Acoustics But Not Comprehension. Journal of Neurophysiology, 104(5):2500–2511. Publisher: American Physiological Society.

Hölle, D. and Bleichner, M. G. (2023). Smartphone-based ear- electroencephalography to study sound processing in everyday life. European Journal of Neuroscience, 58(7):3671–3685. eprint: https://onlinelibrary.wiley.com/doi/pdf/10.1111/ejn.16124.

Hölle, D., Meekes, J., and Bleichner, M. G. (2021). Mobile ear-EEG to study auditory attention in everyday life: Auditory attention in everyday life. Behavior Research Methods, 53(5):2025–2036.

Korte, S., Haupt, T., and Bleichner, M. G. (2025a). EEG Signatures of Auditory Distraction: Neural Responses to Spectral Novelty in Real-World Soundscapes. Pages: 2025.04.14.648656 Section: New Results.

Korte, S., Jaeger, M., Rosenkranz, M., and Bleichner, M. G. (2025b). From beeps to streets: unveiling sensory input and relevance across auditory contexts. Frontiers in Neuroergonomics, 6:1571356. Publisher: Frontiers.

Kriegeskorte, N. and Douglas, P. K. (2019). Interpreting encoding and decoding models. Current Opinion in Neurobiology, 55:167–179.

Ladouce, S., Mustile, M., and Dehais, F. (2021). Capturing cognitive events embedded in the real-world using mobile EEG and Eye-Tracking. Pages: 2021.11.30.470560 Section: New Results.

Lalor, E. C., Power, A. J., Reilly, R. B., and Foxe, J. J. (2009). Resolving Precise Temporal Processing Properties of the Auditory System Using Continuous Stimuli. Journal of Neurophysiology, 102(1):349–359. Publisher: American Physiological Society.

Lanting, C. P., Briley, P. M., Sumner, C. J., and Krumbholz, K. (2013). Mechanisms of adaptation in human auditory cortex. Journal of Neurophysiology, 110(4):973–983. Publisher: American Physiological Society.

Lee, A. K. C., Larson, E., Maddox, R. K., and Shinn-Cunningham, B. G. (2014). Using neuroimaging to understand the cortical mechanisms of auditory selective attention. Hearing Research, 307:111–120.

Lutzenberger, W., Pulvermüller, F., and Birbaumer, N. (1994). Words and pseudowords elicit distinct patterns of 30-Hz EEG responses in humans. Neuroscience Letters, 176(1):115–118.

López-Caballero, F. (2025). N1 facilitation at short Inter-Stimulus-Interval (ISI) occurs under 400 ms and is dependent on ISI from previous sounds: Evidence using an unpredictable auditory stimulation sequence.

López-Caballero, F., Coffman, B., Seebold, D., Teichert, T., and Salisbury, D. F. (2023). Intensity and inter-stimulus-interval effects on human middle- and long-latency auditory evoked potentials in an unpredictable auditory context. Psychophysiology, 60(4):e14217. eprint: https://onlinelibrary.wiley.com/doi/pdf/10.1111/psyp.14217.

May, P. J. C. and Tiitinen, H. (2010). Mismatch negativity (MMN), the deviance- elicited auditory deflection, explained. Psychophysiology, 47(1):66–122. eprint: https://onlinelibrary.wiley.com/doi/pdf/10.1111/j.1469-8986.2009.00856.x.

Mesgarani, N., David, S. V., Fritz, J. B., and Shamma, S. A. (2009). Influence of Context and Behavior on Stimulus Reconstruction From Neural Activity in Primary Auditory Cortex. Journal of Neurophysiology, 102(6):3329–3339. Publisher: American Physiological Society.

Mesik, J. and Wojtczak, M. (2023). The effects of data quantity on performance of temporal response function analyses of natural speech processing. Frontiers in Neuroscience, 16.

Mirkovic, B., Debener, S., Jaeger, M., and De Vos, M. (2015). Decoding the attended speech stream with multi-channel EEG: implications for online, daily- life applications. Journal of Neural Engineering, 12(4):046007. Publisher: IOP Publishing.

Müller, M. (2021). Fundamentals of Music Processing: Using Python and Jupyter Notebooks. Springer International Publishing, Cham.

Näätänen, R. (2001). The perception of speech sounds by the human brain as reflected by the mismatch negativity (MMN) and its magnetic equivalent (MMNm). Psychophysiology, 38(1):1–21.

Näätänen, R. and Picton, T. (1987). The N1 Wave of the Human Electric and Magnetic Response to Sound: A Review and an Analysis of the Component Structure. Psychophysiology, 24(4):375–425. eprint: https://onlinelibrary.wiley.com/doi/pdf/10.1111/j.1469-8986.1987.tb00311.x.

Okamoto, H., Ross, B., Kakigi, R., Kubo, T., and Pantev, C. (2004). N1m recovery from decline after exposure to noise with strong spectral contrasts. Hearing Research, 196(1):77–86.

Rahman, M., Willmore, B. D. B., King, A. J., and Harper, N. S. (2020). Simple transformations capture auditory input to cortex. Proceedings of the National Academy of Sciences, 117(45):28442–28451.

Regev, T. I., Markusfeld, G., Deouell, L. Y., and Nelken, I. (2021). Context Sensitivity across Multiple Time scales with a Flexible Frequency Bandwidth. Cerebral Cortex, 32(1):158–175.

Rosburg, T. and Mager, R. (2021). The reduced auditory evoked potential component N1 after repeated stimulation: Refractoriness hypothesis vs. habituation account. Hearing Research, 400:108140.

Rosburg, T., Weigl, M., and Mager, R. (2022). No evidence for auditory N1 dishabituation in healthy adults after presentation of rare novel distractors. International Journal of Psychophysiology, 174:1–8.

Rosenkranz, M., Cetin, T., Uslar, V. N., and Bleichner, M. G. (2023). Investigating the attentional focus to workplace-related soundscapes in a complex audio-visual-motor task using EEG. Frontiers in Neuroergonomics, 3.

Rosenkranz, M., Haupt, T., Jaeger, M., Uslar, V. N., and Bleichner, M. G. (2024). Using mobile EEG to study auditory work strain during simulated surgical procedures. Scientific Reports, 14(1):24026. Publisher: Nature Publishing Group.

Ruusuvirta, T. (2021). The release from refractoriness hypothesis of N1 of event- related potentials needs reassessment. Hearing Research, 399:107923.

Schutz, M. and Gillard, J. (2020). On the generalization of tones: A detailed exploration of non-speech auditory perception stimuli. Scientific Reports, 10(1):9520. Publisher: Nature Publishing Group.

Shine, J. M., Lewis, L. D., Garrett, D. D., and Hwang, K. (2023). The impact of the human thalamus on brain-wide information processing. Nature Reviews Neuroscience, 24(7):416–430. Publisher: Nature Publishing Group.

Silva, D. M. R., Melges, D. B., and Rothe-Neves, R. (2017). N1 response attenuation and the mismatch negativity (MMN) to within- and across-category phonetic contrasts. Psychophysiology, 54(4):591–600. eprint: https://onlinelibrary.wiley.com/doi/pdf/10.1111/psyp.12824.

Somervail, R., Perovic, S., Bufacchi, R. J., Caminiti, R., and Iannetti, G. D. (2025). A Two-system Theory of Sensory-evoked Brain Responses.

Somervail, R., Zhang, F., Novembre, G., Bufacchi, R. J., Guo, Y., Crepaldi, M., Hu, L., and Iannetti, G. D. (2021). Waves of Change: Brain Sensitivity to Differential, not Absolute, Stimulus Intensity is Conserved Across Humans and Rats. Cerebral Cortex, 31(2):949–960.

Stam, C. J. (2005). Nonlinear dynamical analysis of EEG and MEG: Review of an emerging field. Clinical Neurophysiology, 116(10):2266–2301.

Ulanovsky, N., Las, L., and Nelken, I. (2003). Processing of low-probability sounds by cortical neurons. Nature Neuroscience, 6(4):391–398. Publisher: Nature Publishing Group.

Vallet, W. and van Wassenhove, V. (2023). Can cognitive neuroscience solve the lab-dilemma by going wild? Neuroscience & Biobehavioral Reviews, 155:105463.

Wang, A. L., Mouraux, A., Liang, M., and Iannetti, G. D. (2008). The Enhancement of the N1 Wave Elicited by Sensory Stimuli Presented at Very Short Inter-Stimulus Intervals Is a General Feature across Sensory Systems. PLOS ONE, 3(12):e3929. Publisher: Public Library of Science.

Wang, Y., Tang, Z., Zhang, X., and Yang, L. (2022). Auditory and cross-modal attentional bias toward positive natural sounds: Behavioral and ERP evidence. Frontiers in Human Neuroscience, 16. Publisher: Frontiers.

Willmore, B. D. B. and King, A. J. (2023). Adaptation in auditory processing. Physiological Reviews, 103(2):1025–1058. Publisher: American Physiological Society.

Winkler, I., Debener, S., Müller, K.-R., and Tangermann, M. (2015). On the influence of high-pass filtering on ICA-based artifact reduction in EEG-ERP. In 2015 37th Annual International Conference of the IEEE Engineering in Medicine and Biology Society (EMBC), pages 4101–4105. ISSN: 1558-4615.

Zacharias, N., König, R., and Heil, P. (2012). Stimulation-history effects on the M100 revealed by its differential dependence on the stimulus onset interval. Psychophysiology, 49(7):909–919. eprint: https://onlinelibrary.wiley.com/doi/pdf/10.1111/j.1469-8986.2012.01370.x.

